# Ancestral Genome Reconstruction

**DOI:** 10.64898/2026.04.16.718917

**Authors:** Cléa Siguret, Margaux Olivier, Cécile Huneau, Mamadou Dia Sow, Pierre-Louis Stenger, Christophe Klopp, Marie-Laure Martin, Jean-Philippe Tamby, Peter Civan, Caroline Pont, Olivier Mathieu, Jerome Salse

**Affiliations:** UCA, INRAE, UMR GDEC, 63000, Clermont-Ferrand, France; UCA, CNRS, Inserm, UMR iGReD, 63000, Clermont-Ferrand, France; INRAE GenoToul Bioinformatics, Castanet Tolosan, France; Université Paris-Saclay, CNRS, INRAE, Université Evry, UMR IPS2, 91190, Gif sur Yvette, France; Université Paris-Cité, CNRS, INRAE, UMR IPS2, 91190, Gif sur Yvette, France; Université Paris-Saclay, AgroParisTech, INRAE, UMR MIA Paris-Saclay, 91120, Palaiseau, France

## Abstract

AGR, for Ancestral Genome Reconstruction, is an automatic publicly available and open-source pipeline to infer paleogenomes from modern species genome comparisons exploiting the concept of inter-species chromosomal synteny relationships’ hierarchical clustering that can be used to unveil how ancestral genomes, genes, sequences and functions have been shaped during million years of present-day plant evolution.

## Introduction

Paleogenomics, which consists of reconstructing ancestral genomes of extinct founders by comparing the genomes of modern species, relies on the rationale in which from an ancestral (possibly extinct) genome that evolved into different extant species through speciation and distinct chromosome shuffling events (fusion, fissions, inversions, and translocations), as well as whole genome duplications (WGD), each of the ancestral chromosomes will derive a subset of modern chromosomal regions sharing conserved genes, *i*.*e*. synteny. Following this evolutionary rationale, when reconstructing ancestral genomes *in silico*, comparative genomics of modern genomes should produce genomic fragments showing independent (non-shared) syntenic blocks, referred to as conserved ancestral regions (CARs), which are considered as ancestral chromosomes in the inferred ancestral genomes, *i*.*e*. paleogenomes. Ancestral genomes can then be used as backbones for both fundamental evolutionary studies and applied inter-crops translational research of key agronomical traits (Salse 2023).

## Material and method

Ancestral genomes are reconstructed using an R-based pipeline AGR (for Ancestral Genome Reconstruction) made publicly available. Orthologs (conserved genes between species) and paralogs (duplicated genes within species) are characterized based on the identification of orthogroups (OGs), *i*.*e*. consisting in all modern genes inherited from a single ancestral gene from their last common ancestor (Tatusov et al., 2003; Chen et al., 2006; Trachana et al., 2011; Waterhouse et al., 2013; Powell et al., 2014). OGs are identified based on the all-against-all BLASTp (Altschul et al., 1990) alignment of the selected proteomes and the orthology inference tool Orthofinder (Emms and Kelly, 2015, 2019) using default parameters. The resulted dataset contains both orthologs and paralogs of a given set of compared genomes (Wapinski et al., 2007; Waterhouse et al., 2010; Simola et al., 2013). OGs are then filtered to keep only conserved gene families (or orthologs) between at least two different species. AGR then reconstruct automatically ancestral genomes at the basis of the considered genomes from a conserved genes’ matrix either obtained from the previous Orthofinder inference, or alternatively from any other tool and methodology users may favor in defining conserved genes between the investigated species, following five steps. **Step #1** (named ‘Matrix Design’) consists in selecting OGs that have, for each selected species, a number of genes equal to (or less than) the expected number of WGD events. This step generates a matrix where OGs are arranged in rows while the chromosome numbers for each species are represented in columns. To differentiate chromosome names between different species, we included the species acronym as prefix of the chromosome. At this stage, the resulting matrix contains both paralogous and orthologous genes between the investigated species. We kept only OGs having at least a number of genes equal to or less than the expected number of duplication events (WGDs). This step required three files: orthogroup, chr and gff files. **Step #2** (named ‘Chromosome-Chromosome relations & Clusters QC’, *i*.*e*. Quality Control) consists in computing the number of k clusters defining the k ancestral blocks (*i*.*e*. the number of ancestral chromosomes). To infer the number of ancestral blocks, the AGR pipeline performs chromosome-to-chromosome hierarchical clustering using Pearson correlation distance and Ward linkage. The obtained graph shows a dendrogram of chromosome-chromosome relationships and a heatmap of the intensity of the synteny relationships between modern chromosomes. We then developed an automatic method to identify the optimal number of k clusters (*i*.*e*. number of ancestral blocks), independently of the orthologous signal strength. We favored the use of height distances directly computed from the dendrogram and applied a twisted Cattell’s rule (1966). Thus, delta (difference in height distance between the clusters) is used for defining optimal k (the highest delta). The quality of k clusters or blocks is then assessed using silhouette width (Rousseeuw, 1987) and Dunn index (Ben Ncir et al., 2021). For both indices, observations around 1 mean high consistency of the chromosome-chromosome clustering based on modern genome synteny relationships. For silhouette index, a value around 0 means that the associated modern chromosomes are placed in the wrong cluster (*i*.*e* ancestral chromosome or CAR). DUNN index (like silhouette index) tests the compactness (small variance between members of the same cluster) and the well separation of clusters (maximum difference between sister clusters). Hence, the two indexes allow to identify whether chromosomes share synteny between two or more ancestral blocks, probably due to past fission/fusion events, classically observed during species evolution. **Step #3** (named ‘Orthogroup-Chromosome relations & CARs definition’). Once the k clusters (number of ancestral blocks) have been validated, the resulting ordered dendrogram and the filtered matrix are used to assess the number of orthogroups in each defined ancestral block by performing Orthogroup-to-Chromosome clustering, where chromosome’s order is fixed according to the previously defined (k) order in step 2. Using the row dendrogram (orthogroups), we identify the number of optimal clusters named kog (group of OGs) like in step 2. At this stage, kog (number of orthogroup clusters) can be equal to or greater than k (number of chromosome clusters). **Step #4** (named ‘Iterative scenario & build pre-ancestor’). When kog > k, *i*.*e*. the number of orthogroup clusters is higher than the expected ancestral blocks, we implemented an iterative scenario that identifies orthogroup clusters to be merged to reach the number of ancestral blocks. This step first identifies theoretical clusters that are prone to fuse (*e*.*g*. clusters 1 & 2, 2 & 3, 3 & 4, etc.) while the fusion of non-successive clusters (like 1 & 3, 2 & 5, etc.) are scored with high penalty. For each identified theoretical fusion, we then compute the number of modern chromosome fragments that would be merged in these scenarios. AGR then selects automatically the inferred ancestor based the scenario that minimize the number of rearrangements in modern species since their common ancestor. For this, different metrics are defined: (*i*) “fusion_prob”: correspond to the theoretical/possible clusters fusions that match the row (orthogroup) dendrogram nodes (TRUE or FALSE); (*ii*) “strength”: if the fusion of two branches (or clusters) is directly feasible without requiring others nodes, then ‘strong’ and if not ‘weak’ are mentioned; (*iii*) “total_fusion”: correspond to the number of modern chromosome fragments that will be merged (into a single fragment) if the corresponding clusters are merged; (*iv*) “merge_height”: correspond to the height of the nodes at each fusion branch node, with the lowest prioritized. From these metrics, the optimal cluster merging is scored with “fusion_prob”=TRUE & “strength”= strong (maximizing the number of “total_fusion” and lowering “merge_height”). This allows to infer conserved ancestral regions (CARs) and to build the most relevant ancestors. For each CAR in the ancestor, the gene order along ancestral chromosomes is determined by using as a reference the modern chromosome with the highest number of genes in the reconstructed CAR. At this step, a quality control of the ancestor can be performed using automatic dotplots and painting graphs of modern chromosomes with ancestral chromosome colors to highlight karyotype evolution. **Step #5** (named ‘Genes’ Enrichment & build final ancestor’) consists in recovering the final number of ancestral protogenes in each ancestral block. Thus, all orthogroups with genes conserved between all species corresponding to the speciation event where the ancestor is reconstructed are added. Inferred CARs are finally validated through the detection of syntenic regions with dotplots, showing that all modern chromosomes in CARs share orthologous and paralogous relationships with the total absence of any syntenic relationship between modern chromosomes of different species inherited from different CARs, the meeting the evolutionary rationale in which from an ancestral genome that evolved into different extant species through speciation, chromosome shuffling events and whole genome duplications, each of the ancestral chromosomes derive a subset of modern chromosomal regions sharing conserved syntenic genes.

## Result

We developed inter-species chromosomal synteny relationships’ hierarchical clustering method, AGR (Ancestral Genome Reconstruction), to infer ancestral chromosome (protochromosome) structure and ancestral gene (protogene) content, from the comparison of modern genomes, following five steps: Step #1 (‘Matrix Design’), Step #2 (‘Chromosome-Chromosome relations & Clusters QC’); Step #3 (‘Orthogroup-Chromosome relations & CARs definition’), Step #4 (‘Iterative scenario & build pre-ancestor’), Step #5 (‘Genes’ Enrichment & build final ancestor’), see Figure below. AGR takes as input a curated orthogroup matrix in which copy numbers are consistent with known whole-genome duplications, and uses both chromosome–chromosome and chromosome-orthogroup hierarchical clustering strategies to define protochromosomes, while providing quantitative indices of clustering quality (silhouette width and Dunn index) to explicitly score the robustness of each inferred ancestors. A key feature of AGR is an iterative merging procedure that reconciles orthogroup clusters with the expected number of ancestral blocks; by prioritizing evolutionary principles established since more than a decade ago, in favoring (1) fusions events in evolution (in the transition from the ancestor to the modern species) known to ensure the maintenance of the centromere-telomere polarity of chromosome arms in the derived dicentric (fused) chromosome with one of the two centromeres inactivated through pericentric inversions (Murat et al. 2010, 2015), and (2) an evolutionary scenario taking into account the fewest number of genomic rearrangements (including inversions, deletions, fusions, fissions and translocations) that may have occurred between the ancestors and modern genomes. Protogene order along each protochromosome is then relying on the modern chromosomes sharing the highest number of conserved genes, and the ancestral gene repertoire is enriched by adding genes conserved at the focal speciation node, before validating the final scenario through modern *vs* ancestor genomes dotplot comparison to ensure that each ancestral chromosome integrates all syntenic segments from extant genomes without cross-contamination (*i*.*e* no shared synteny between modern chromosomes deriving from distinct inferred ancestral chromosomes). AGR addresses a long-standing limitation of ancestral reconstruction in plants, namely, the difficulty of balancing synteny signal, duplication noise, and rearrangements across deeply diverged lineages, and now delivers statistically supported ancestors and evolutionary scenarios that can be directly compared across studies.

**Figure:**
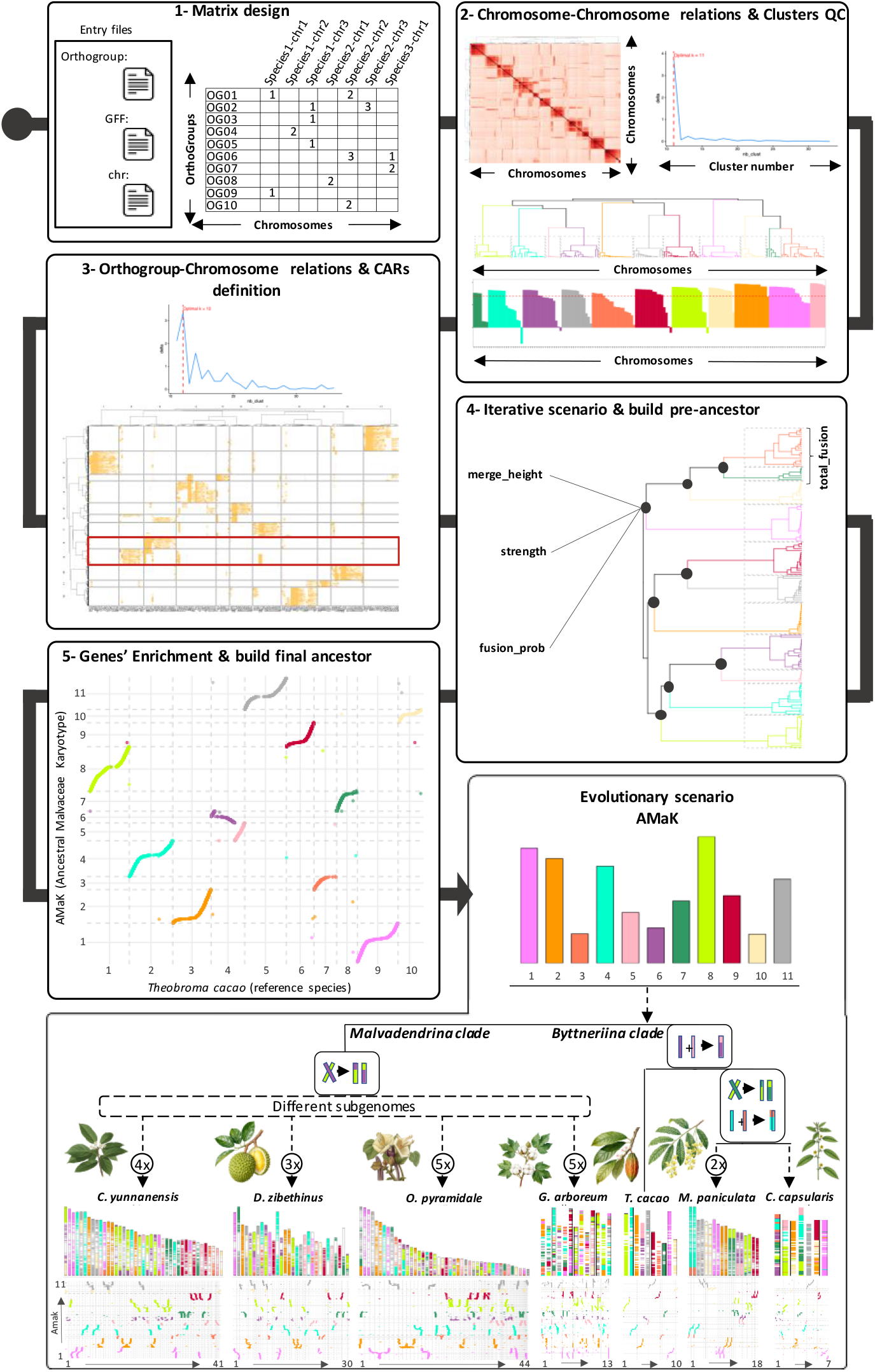
Ancestral genome reconstruction (AGR) tool. Reconstruction of ancestral karyotypes and evolutionary scenarios, using the Malvaceae family as a tutorial case example. The figure illustrates the inferred chromosomal relationships, clustering analyses, and evolutionary events leading to the diversification of modern Malvaceae genomes. Panels (i–ii) show the hierarchical clustering and delta plot used to determine the optimal number of chromosome clusters (k = 11–12). Panel (iii) displays the cluster dendrogram defining chromosomal affinities (synteny) among the considered species. Panel (iv) presents a silhouette plot validating cluster robustness. Panel (v) summarizes the iterative reconstruction of protochromosomal regions and their hierarchical fusion events to build the pre-ancestral karyotype (pre-AMaK). Panel (vi) depicts the final ancestral Malvaceae karyotype (AMaK), derived from comparative synteny analyses among seven representative modern species (Theobroma cacao, Microcos paniculata, Corchorus capsularis, Craigia yunnanensis, Durio zibethinus, Ochroma pyramidale, and Gossypium arboretum). The lower schematic illustration represents an evolutionary scenario separating the Byttneriina and Malvadenrina clades, with successive polyploidization events (referred to as 2× to 5×) and (sub-)genome differentiation across lineages. Colored bars and synteny links indicate conserved chromosomal segments and genome rearrangements that shaped the diversification of modern Malvaceae from AMaK.

We illustrate AGR on a case study example providing a detailed step-by-step tutorial in using AGR for reconstructing ancestral genomes of the well-studied *Malvaceae* family to infer AGR-based AMaK (for Ancestral *Malvaceae* Karyotye) published recently (Sun et al., 2024, Shao et al. 2024 and Zhang et al. 2025) from the comparison of modern genomes across the *Byttneriina* (*Gossypium arboreum Theobroma cacao, Microcos paniculata*, and *Corchorus capsularis)* and *Malvadenrina* (*Craiga yunnanensis, Durio zibethinus, Ochroma pyramidale)* clades. The reconstruction integrated orthogroup assignments, gene annotations, and chromosomal information from seven representative modern species across the *Malvaceae* family. This reconstruction seeks to elucidate the evolutionary history and principal genomic architecture of this plant family, as well as explained in more details the relevance of our pipeline. The first step involves the careful selection of extant species that collectively capture the phylogenetic breadth of *Malvaceae*. Species are chosen based on two major criteria: (1) their position within key evolutionary branches to ensure coverage of all major lineages, and (2) their genome architecture, favoring those with limited recent whole-genome duplication and clear structural diversification. This deliberate choice ensures that the evolutionary signals for reconstructing the ancestral state are as robust and representative as possible. Genomic resources collated for each species include: annotated gene models, chromosomal structure data, and gene families (orthogroups). Central to ancestral reconstruction is the comparative analysis of gene families across the selected genomes. Genes are organized into orthogroups, which represent collections of homologous sequences descended from a common ancestor. The workflow emphasizes orthogroups with conserved copy number and chromosomal location across species, as these are most informative for inferring ancestral relationships. Rather than simply cataloging homologs, the approach critically evaluates gene duplication, loss, and divergence events, considering lineage-specific genome dynamics. The aim is to remove confounding effects from recent duplications and concentrate on gene families whose evolutionary histories are likely shared with the ancestor. To reflect how chromosomes have been conserved, reordered, or split during evolution, a matrix correlating the presence of orthogroups across chromosomes is constructed. This matrix enables the direct identification of chromosomes with shared ancestry, those which preferentially retain the same collections of orthogroups. Hierarchical clustering of this matrix provides an objective means to infer ancestral chromosomal blocks, termed Contiguous Ancestral Regions (CARs). The rationale is that chromosomes clustering together by orthogroup content likely reflect conserved blocks from the ancestral karyotype. The number of ancestral blocks is carefully estimated by considering both the chromosomal counts in modern species and the extent of whole genome duplication events. Statistical validation, including silhouette and Dunn indices, assesses the robustness of the inferred clusters and the biological plausibility of the ancestral chromosome number. Having identified initial clusters of orthogroups and chromosomal regions, the workflow seeks to reconcile discrepancies between gene-based and chromosome-based groupings. Conflicts are resolved through examination of evolutionary events such as chromosomal fusions and fissions, employing the principle of parsimony to favor the scenario requiring the least number of structural rearrangements between ancestor and descendants. The merging (fusion) of ancestral blocks is guided by evidence of conserved co-occurrence in modern species and by their hierarchical proximity in the clustering dendrogram. This conceptual step ensures that the final set of ancestral regions reflects both evolutionary logic and empirical patterns observed across the investigated species family. With ancestral blocks defined, the workflow models the organization of the ancestral genome by projecting syntenic relationships from modern species onto the ancestor. For each CAR, representative sequences from extant chromosomes showing strongest homology are selected to guide the ordering of genes. This step acknowledges that while gene identity can often be robustly traced, gene order may have undergone rearrangements, and thus, projecting the most conserved order provides the best available reconstruction. Recognizing that gene assignments may be affected by artifacts, such as incomplete lineage sorting or gene transfer, the method implements rigorous filtering based on synteny the conservation of sequential gene order across species. Only genes sustained within conserved syntenic blocks across multiple taxa are retained in the ancestral model, ensuring that the reconstruction is relying on strong evolutionary evidence. This step filters out anomalous gene associations, thus strengthening the biological authenticity of the inferred ancestor. To avoid underrepresentation of the ancestral gene repertoire, the model is enriched with additional gene families conserved between a subset of the analyzed species. By incorporating orthogroups present in at least 10% of *Malvaceae* genomes, the workflow accounts for lineage-specific gene losses and ensures a more complete repertoire of ancestral gene content. New genes are integrated in respect of syntenic relationships and known evolutionary constraints, refining both completeness and accuracy of the reconstructed ancestor. The workflow concludes with extensive validation through chromosomal painting and synteny dotplots. Chromosome painting color-codes modern chromosomes according to their inferred ancestral blocks, revealing patterns of ancestral block retention and rearrangement in the modern genomes. Synteny dotplots visualize the correspondence between ancestral and modern genomes, then highlighting collinearity, breakpoints, and copy number variations that narrate the evolutionary trajectory from ancestral to modern karyotypes. These visual validations reinforce the plausibility of the reconstructed ancestor and illustrate the evolutionary logic connecting extant genomes to their inferred ancestors.

Here, each methodological choice is motivated by the aim to accurately recapitulate the most parsimonious and biologically faithful ancestral genomes, offering a robust foundation for subsequent evolutionary analysis in *Malvaceae*, with the (i) determination of the optimal number of clusters (k = 11) based on the delta value of the clustering index, (ii) hierarchical clustering dendrogram displaying the 11 identified clusters (each represented by a different color), (iii) silhouette plot illustrating the consistency and robustness of clustering assignments, with an average silhouette width of 0.59, (iv) summary statistics of cluster characteristics, including cluster size and average silhouette width for each group, (v) iterative reconstruction of protochromosomal regions from hierarchical cluster fusion events to build the ancestral karyotype (AMaK), (vi) distribution of clusters (protochromosomes) across inferred ancestral (y-axis) and modern (x-axis) genomes through two-dimensional ordination plots (synteny plots), with colored blocks and synteny links depicting conserved genomic regions and highlighting the complex history of chromosomal rearrangements and genome duplication events across *Malvaceae* : a shared reciprocal translocation at the basis of the *Malvadenrina* and a shared ancestral chromosome fusion within the *Byttneriina* (followed by species specific reciprocal translocation and chromosome fusion for *Microcos paniculate* and *Corchorus capsularis*), polyploidization events referred to as 2× to 5× for *Microcos paniculata, Craigia yunnanensis, Durio zibethinus, Ochroma pyramidale, Gossypium arboretum*, as well as lineage-, genome- and even subgenome-specific rearrangements as detailed in https://forge.inrae.fr/umr-gdec/ancestor-malvaceae-reconstruction.

## Conclusion

AGR turns ancestral genome reconstruction from an opaque, method-driven exercise into a transparent, testable framework to propose quality controlled ancestral genomes either at the basis of major plant clades (angiosperms, monocots, dicots) or botanical families (legumes, cereals, *Triticeae*…), Murat et al. 2017. Ancestral genomes can be used then to unveil how ancestral genomes, genes, sequences and functions have been shaped during millions of years of plant genome evolution and finally to improve the efficient transfer of innovation between species for crops’ improvement, as detailed in the ***full version of this article currently under peer review in scientific journal***.

## Acknowledgments

We are grateful to the genotoul bioinformatics platform Toulouse Occitanie (Bioinfo Genotoul, https://doi.org/10.15454/1.5572369328961167E12) and the Mesocentre Clermont Auvergne bioinformatics platform (https://doi.org/10.18145/aubi) for providing computing and storage resources. This work was supported by the ‘Région Auvergne-Rhône-Alpes’ and FEDER funding (SRESRI 2015, PaleoLAB), Institut Carnot Plant2Pro (projects SyntenyViewer https://plant2pro.fr/wp-content/uploads/SYNTENYVIEWER.pdf and OrthoBreeding https://plant2pro.fr/wp-content/uploads/Orthobreeding-2023.pdf), the ANR (projects PAGE https://anr.fr/Project-ANR-11-BSV6-0008, AkaeoAG, ANR-20-CE27-0013 https://anr.fr/Projet-ANR-20-CE27-0013), The ISITE CAP2025 (#00002146 SRESRI ‘Pack Ambition Recherche Project’ TransBlé https://cap2025.fr/fr/news/actualites-en-cours/actualites-passees/archives-2018/resultats-de-lami-pack-ambition-recherche-2018), and PEPR ‘Agroecology and Numeric’ (Flagship AgroDIV, 22-PEAE-0005 https://anr.fr/ProjetIA-22-PEAE-0005).

## Author contributions

C. Siguret, M. Olivier, C. Huneau, M. D. Sow, P.-L. Stenger performed the analysis, delivered the associated illustration and provided their contribution to the manuscript; C. Klopp, M.L. Martin, J.P. Tamby, P. Civan, C. Pont, contributed with key complementary expertise; O. Mathieu and J. Salse supervised the analysis and the article preparation.

## Data and Material Availability

Ancestral genome reconstruction (AGR) pipeline in R is made publicly available with a tutorial example on the *Malvaceae* family: https://forge.inrae.fr/umr-gdec/ancestor-malvaceae-reconstruction (*first public release on October 2025, 29*^*th*^).

